# Real-time identification of epistatic interactions in SARS-CoV-2 from large genome collections

**DOI:** 10.1101/2023.08.22.554253

**Authors:** Gabriel Innocenti, Marco Galardini

## Abstract

The emergence and rapid spread of the SARS-CoV-2 virus has highlighted the importance of genomic epidemiology in understanding the evolution of pathogens and for guiding public health interventions. In particular, the Omicron variant underscored the role of epistasis in the evolution of lineages with both higher infectivity and immune escape, and therefore the necessity to update surveillance pipelines to detect them as soon as they emerge. In this study we applied a method based on mutual information (MI) between positions in a multiple sequence alignment (MSA), which is capable of scaling up to millions of samples. We showed how it could reliably predict known experimentally validated epistatic interactions, even when using as little as 10,000 sequences, which opens the possibility of making it a near real-time prediction system. We tested this possibility by modifying the method to account for sample collection date and applied it retrospectively to MSAs for each month between March 2020 and March 2023. We could detect a cornerstone epistatic interaction in the Spike protein between codons 498 and 501 as soon as 6 samples with a double mutation were present in the dataset, thus demonstrating the method’s sensitivity. Lastly we provide examples of predicted interactions between genes, which are harder to test experimentally and therefore more likely to be overlooked. This method could become part of continuous surveillance systems tracking present and future pathogen outbreaks.

## Introduction

The COVID-19 severe respiratory syndrome is caused by the SARS-CoV-2 virus, which emerged in late 2019 in China and quickly escalated to a pandemic. Since then, the virus has differentiated into a number of lineages, almost all of which have taken over the whole population in successive sweeps reminiscent of clonal interference^1^. These successful lineages - designated as Variants Of Concern (VOC) - have acquired a number of mutations that have increased its ability to infect new hosts and escape infection and vaccine induced immunity^2–9^. The unprecedented efforts in the genomic epidemiology of the virus leading to millions of whole sequences being deposited in public and restricted-access databases has accelerated the pace by which emerging lineages are tracked and the impact of genetic mutations are estimated^1,10–12^. Together with expedited experimental measurements of the impact of single mutations it has been possible to obtain estimates of the fitness advantage of emerging variants with as little delay as a few weeks^13–15^. Given the successes of SARS-CoV-2 genome sequencing in tracking the emergence and spread of variants and to develop groundbreaking vaccines^16^, it is hard to imagine it not becoming a routine tool in handling current and future epidemics^17^.

Which other information of relevance can be obtained from large genome sequencing datasets? A growing body of evidence from population genetics and evolutionary studies indicates that the impact of individual genetic variants can be modulated by the presence of other variants in other sites, a phenomenon known as epistasis or genetic background effect^18^. In practice this means that the measured impact of a genetic variant may be very different when the same variant is present in a different genetic background, which in turn makes sequence and phenotypic evolution unpredictable at medium to long timescales^19^. In the context of SARS-CoV-2 evolution, epistasis might have implications for genomic surveillance of lineages; assumptions about the fitness advantage conferred by a mutation, be it through increased transmissibility or immune escape, might be invalidated and lead to erroneous applications of public health measures or vaccine updates. At a smaller scale, this possibility has been experimentally confirmed for the receptor binding domain (RBD) of the SARS-CoV-2 spike protein, for which mutations have a different impact on antibody escape depending on the overall genetic background (*i.e.* different VOCs)^20,21^. Detecting potential epistatic interactions between mutating sites in the SARS-CoV-2 genomes could be used as a sign that the fitness effect of a particular mutation in a particular genetic background might not generalize in another. Furthermore, positions participating in an epistatic interaction might indicate that a particular fitness “peak” can only be reached through another neutral or slightly deleterious mutation^22^. Predicting whether seemingly neutral mutations are enabling further ones that affect transmissibility or immune escape would therefore be a valuable epidemiological tool. Lastly, from the perspective of genomic surveillance, being able to quickly identify sites that participate in an epistatic interaction would help identify virus’ variants that could have a fitness advantage and that could therefore quickly spread.

In order to make the estimation of epistatic interactions useful in the context of a rapidly unfolding pandemic, the method needs to have an appropriate combination of speed and precision/sensitivity. Ideally it would require modest computational resources and would run in near real-time, so that it could be implemented as part of existing automated genomic epidemiology tools that feed on sequence repositories^1,10,23^. Some of the computational approaches that are able to estimate epistatic interactions would not be suitable for this task for resource considerations, such as pseudolikelihood or phylogenetic methods^24–30^. Aggressive subsampling would make these methods applicable, but could theoretically reduce their ability to quickly identify new interactions and variants that are starting to emerge.

Here, we explore the usefulness of a method based on the detection of mutual information (MI) between sites in a multiple sequence alignment^31^ as an indirect estimate of mutational epistasis, which is able to handle alignments with more than 10^6^ samples. We applied it to all high-quality publicly available SARS-CoV-2 sequences as of April 14th 2023 (N=6,644,032). The method detected 474 putative epistatic interactions between different positions, 222 of which with high mutual information. We validated the highest scoring hits using a list of known Mutation Of Interest/Concern (MUI and MOC)^32^, data from deep mutational scans^20,21^, and different epistasis models^24–26^. We further adapted the method to account for the “age” of each sequence, leveraging the metadata associated with each sequence, and thus showing how interactions gain/loss varies over time and variant emergence, and how this could be used as a near real-time genomic surveillance system. To this extent we demonstrated how our method is highly sensitive, being able to identify a known epistatic interaction in the Omicron variant with as little as six sequences. Taken together, these results offer yet another vision of the future of pathogen genomic epidemiology, in which the subtle complexities of evolution of biological sequences are taken into account.

## Results

### Mutual information methods can estimate epistatic interactions in a large scale genomic dataset

This analysis was possible thanks to the data available on the GISAID database, from which we downloaded the SARS-CoV2 multialignment file and its relative phylogenetic tree. After 2 steps of filtering, followed by trimming and deduplication, a phylogenetic weighting strategy was applied to each sequence in the alignment in order to account for population structure. Finally the spydrpick algorithm^31^ was run to compute mutual information (MI) between every pair of positions in the genome (pipeline shown in **Figure 1A**). Each position pair was then assigned to an outlier level O (O1, O2, O3, O4) according to their score, with O4 values indicating the strongest predicted interactions (see **Methods**).

**Figure 1.**
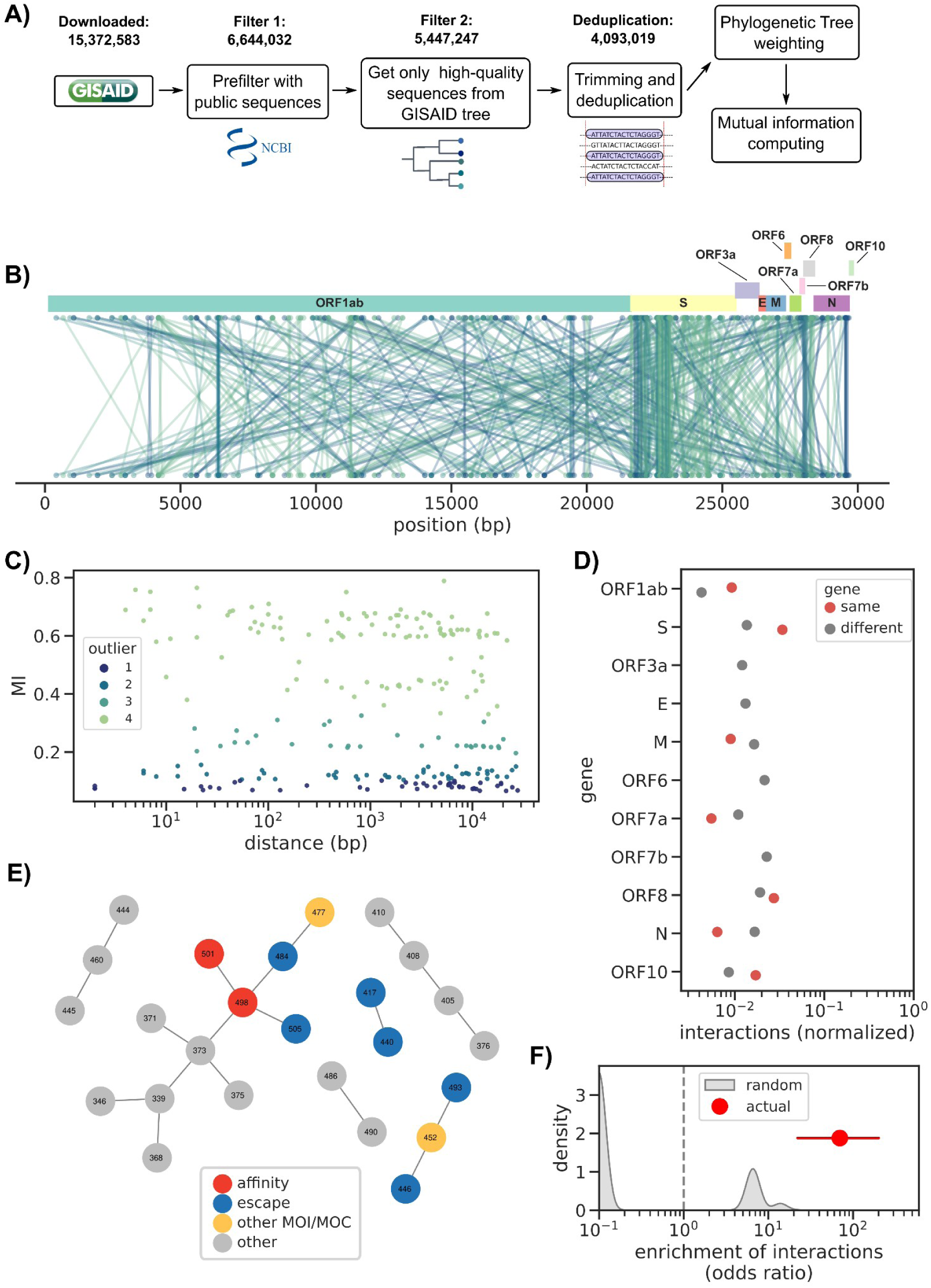
Estimation of epistatic interactions from >4M SARS-CoV-2 sequences. A) Analysis workflow. B) Overview of the estimated epistatic interactions across the whole SARS-CoV-2 genome; each interaction is colored according to its outlier level (O1 to O4), using the same color scale as panel C. C) Distance distribution between all interactions. D) Proportion of interactions within and between genes, normalized by the nucleotide length of each focal gene. E) Interaction network for the RBD region of the Spike gene; amino acid positions are colored according to publicly available annotations. F) Enrichment of interactions between positions known to epistatically interact (red dot) versus a series (N=1,000) of random RBD networks with the same number of interactions as the real one (gray distribution). The red horizontal line indicates the 95% confidence interval.

The unfiltered dataset counted 15,372,583 sequences (up to 14th April 2023). From these we picked the public sequences (*i.e.* also available in the NCBI’s Genbank database, N=6,644,032) which were then filtered by using only the high quality sequences present in the GISAID SARS-CoV2 phylogenetic tree and by removing exact duplicates (N=4,093,019).

We obtained 474 interactions (between 247 unique positions) with many interactions both within and between different genes (**Figure 1B**, **Supplementary Table 1**). Most of the observed interactions were concerning the ORF1ab and Spike gene, with 179 and 185 interactions each, respectively. The interactions were assigned to 4 “outliers levels” (O1 to O4) based on their MI value (**Methods**), with an overrepresentation of O4 interactions (NO4=222) compared to the others which showed similar frequencies (NO1=92, NO2=96, NO3=64). However, they showed similar distribution in function of the distance between the interacting genome positions (**Figure 1C**).

Another important aspect that we considered was if the interactions fell within the same gene or between different ones: ORF3a, the Envelope gene (E), ORF6 and ORF7b presented only interactions between different genes, while the remaining had both kinds. In particular, Spike, ORF1ab and the Nucleocapsid gene (N) presented the highest ratio between different/same gene interactions using normalized counts (**Figure 1D**), showing more interactions in the same gene compared to the others.

To further validate our predictions, we labeled positions in the Spike RBD as either affinity mutations^21^, escape mutations^20^ and other MOI/MOC^32^. These positions have been shown to engage in epistatic interactions resulting in increased viral fitness through higher infectivity or immune escape. The RBD interaction’s network we obtained from our predictions (shown in **Figure 1E**) captured all 3 kinds of mutations. Among them, the interaction between affinity mutations 501 and 498 was observed, but also between escape mutations and other mutations of interest including 484, 505, 446, 477. In addition, other positions with unknown significance (*e.g.* 373, 390, 486, 376) were highlighted by our method, and would need a deeper investigation to understand if they engage in actual epistatic interactions or if they are the false positives. We verified that the reconstructed interaction network was significantly different than a random one via a permutation test, which showed a strong enrichment in known epistatic interactions in the RBD region of the Spike gene (odds-ratio 69.9 [22.0-202.1], **Figure 1F**). We observed an even stronger enrichment (odds-ratio 104.8 [42.9-336.0]) when restricting interactions with outlier level O3 and O4. We further used the outlier thresholds O3 and O4 to build a binary classifier for known RBD epistatic interactions (**Methods**), which had a specificity of 0.33 [0.17-0.71] and 0.50 [0.17-0.71], and a sensitivity of 1 and 0.71, respectively. Lastly, we used the pairwise epistasis coefficients for 15 BA.1 mutations^21^ and measured the ability to recover those with a value over 0.4 using O3 and O4 interactions, for which we recorded a specificity of 0.93 [0.86-0.97] and sensitivity of 0.4 for both thresholds. These results showed how this method was able to pick many experimentally verified interactions and possible new ones.

### Influence of dataset size on the estimation of epistatic interactions

Even though our implementation of the spydrpick algorithm can be run in reasonable time in a high-performance cluster (*i.e.* ∼36 hours, each job requiring > 150Gb of RAM), we reasoned that for it to be of real use it would need to work with a leaner dataset. We therefore created 4 smaller datasets with orders of magnitude less sequences than the complete >4M dataset (N=1,000, 10,000, 100,000, and 1,000,000, randomly chosen). We observed that the number of predicted interactions was larger with smaller datasets, and that the numbers became comparable with the complete dataset when at least 100,000 sequences were considered (Figure 2A, **Supplementary Material 1**). Despite the large difference in predicted interactions, we observed a strong correlation (*r* > 0.95) on MI values for those interactions found both in a reduced subset and the complete dataset (Figure 2B-E). The overlap became larger with subset size, from 57.8% of the 474 interactions from the complete dataset predicted in the N=1,000 subset to 92.4% in the N=1,000,000 subset. We also observed a stronger enrichment in known epistatic interactions in the Spike RBD with increasing sample size, with comparable strength as the complete dataset when using at least 100,000 sequences (odds-ratio > 10 for all subsets, Figure 2F-I). When reducing the interactions to outlier levels O3 and O4 we observed a consistently high level of enrichment across all subsets (odds-ratios > 100), which could be explained by the high number of predicted interactions with outlier level O1 and O2 in the smaller 1,000 and 10,000 datasets (**Supplementary Figure 1**). We then used the O3 outlier threshold to build a binary classifier for RBD known epistatic interactions, which yielded the following specificity/sensitivity values: 0.97 [0.95-0.99] / 0.19 [0.09-0.28] (n=1,000), 0.80 [0.68-0.96] / 1.00 (N=10,000), 0.46 [0.28-0.84] / 1.00 (N=100,000), and 0.38 [0.21-0.74] / 1.00 (N=1,000,000). When predicting pairwise interactions with coefficient > 0.4 using O3 interactions we instead recorded the following specificity/sensitivity values: 0.93 [0.86-0.97] / 0.40 (n=1,000), 0.94 [0.87-0.98] / 0.40 [0.00-1.00] (N=10,000), 0.94 [0.88-0.98] / 0.40 (N=100,000), and 0.93 [0.87-0.97] / 0.40 (N=1,000,000). We therefore estimate that mutual information based estimation of epistatic interactions in SARS-CoV-2 can be performed with datasets with size between 10,000 and 100,000 sequences.

**Figure 2.**
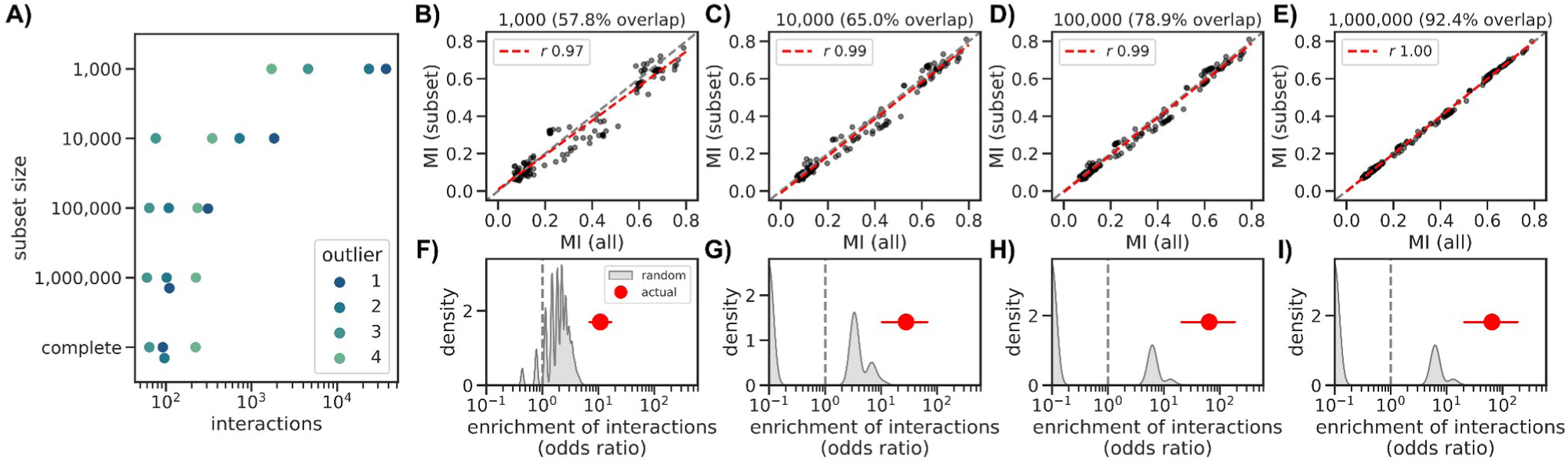
Influence of dataset size on mutual information estimation. A) Number of interactions across datasets, divided by outlier level; the “complete” dataset refers to the one computed using 4,093,019 sequences. B-E) Linear regression analysis of mutual information values in the complete dataset (x-axis) versus the same interactions in the 1,000 (B), 10,000 (C), 100,000 (D), and 1,000,000 (E) subsets. The dashed red line shows the linear regression, and the corresponding r-value is indicated in the panel legend. F-I) Enrichment of interactions between positions known to epistatically interact (red dots) versus a series (N=1,000) of random RBD networks with the same number of interactions as the real one (gray distribution). Subsets are the same as the panel directly above: 1,000 (F), 10,000 (G), 100,000 (H), and 1,000,000 (I). The red lines indicate the 95% confidence interval.

### Real-time estimation of epistatic interactions

The unprecedented genomic epidemiology effort during the SARS-CoV-2 pandemic offers the opportunity to test the usefulness of computational methods to quickly identify emerging pathogen variants and mutations of concern^15^. In the same spirit we sought to understand if we could apply the mutual information methodology described here to develop a near real-time system to discover emerging epistatic interactions. We therefore used the sample collection date associated with each viral sequence to divide the complete dataset into 37 subsets, one for each month from March 2020 up to March 2023, and selecting up to 2,500 sequences for each month. We then computed the mutual information for each month using the cumulative sequences up until the focal month (Figure 3A). In order to highlight emerging interactions and remove earlier ones we introduced a further weight to each sequence based on its distance in time from the focal month. That is, we placed lower importance to older sequences, halving their weight at around six months (120 days), using a hill function (**Methods**, **Supplementary Figure 2**). We reasoned that this would allow the method to discard older interactions and focus on emerging ones.

**Figure 3.**
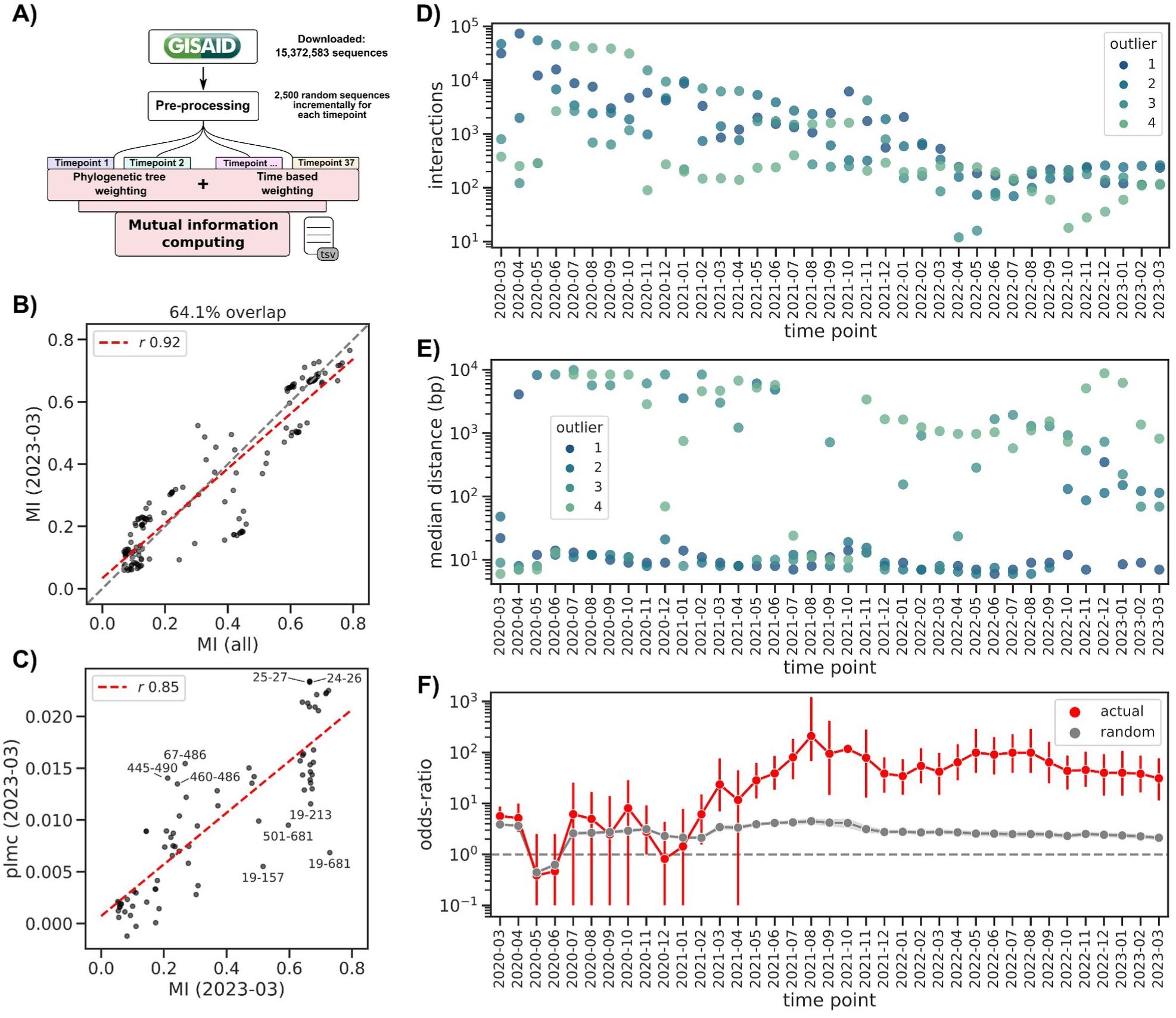
Time based mutual information estimation. A) Analysis workflow. B) Comparison between MI values in the total dataset versus those in the last subset (March 2023, without the time weighting correction). C) Comparison of MI values in the Spike gene between the March 2023 subset and the output of the plmc pseudo-likelihood method; the 10 interactions with the highest absolute residuals with respect to the linear regression are labeled with the amino acid position in the Spike gene. D) Number of interactions for each outlier category across all the time subsets. E) Median distance between interacting positions, divided by outlier level and across all the time subsets. F) Enrichment of interactions between positions known to epistatically interact (red dots) versus a series (N=1,000) of random RBD networks with the same number of interactions as the real one (gray line, shaded area represents the standard deviation, vertical red lines indicate the 95% confidence interval).

We first verified that the final subset yielded comparable results as larger datasets and as a partially orthogonal method^24–26^. For this we used the last subset (March 2023, N=84,923) and avoided the time weighting step to allow for a fair comparison. We observed that MI values had a high correlation with the complete dataset, with 64.1% of the interactions found in the complete dataset recovered in the final subset and with a linear regression r-value of 0.92 (Figure 3B, **Supplementary Table 2**). We also compared the correlation between the MI values against estimates using a pseudo-likelihood method^24–26^ (**Methods**). For this analysis we focused only on the Spike gene as the plmc implementation does not scale well to genome-scale nucleotide alignments. We again observed a very high correlation (linear regression *r*-value of 0.85, Figure 3C, **Supplementary Material 2**). We singled out the 10 interactions with the highest residual: two of them (amino acid positions 460/486 and 445/490) belonged to the RBD region of the spike and had a higher interaction score when using the pseudolikelihood method. Conversely the interaction between the mutations of concern 501/681 was scored higher when using the MI based method.

We next computed MI values for all subsets with the time correction; the number of estimated interactions decreased steadily over time, from a total of 79,586 for the March 2020 subset to 730 in the final March 2023 subset. The number of O4 interactions was more stable over time, especially starting from 2021: the median number of O4 interactions in 2020 was 1,513 versus 196 in the 2021 to 2023 period (Figure 3D, **Supplementary Material 3**). The median distance between interacting nucleotide positions was more stable over time, with O3/O4 interactions being more distant from each other than lower confidence ones: the median distance was 284 and 1,232 bases over all subsets, respectively, while O1/O2 interactions’ median distance was 9 and 12 bases, respectively (Figure 3E). A lower distance between interactions could be a sign of a lower confidence interaction, at least in the general case.

In line with the fact that variants of concern with both higher infectivity and immune escape capabilities did not emerge before the spring of 2021 (*e.g.* the Delta variant), we did not observe an appreciably high enrichment of known interactions between affinity and escape Spike mutations before at least July 2021 (odds-ratio 80.2 [29.5-184.2]), and we observed a consistently high enrichment level until the last time point (odds-ratio > 30, Figure 3F, **Supplementary Table 3**). We measured the ability of this method to recover known RBD interactions by assessing the F1 score, specificity and sensitivity of binary classifiers built using the outlier thresholds (**Supplementary Figure 3**). The O4 binary classifier had the highest F1-score in the December 2012 dataset (value 0.8 [0.61-0.91]), consistent with the emergence of the omicron variant, and no predictive power before then. When we used the predicted RBD interaction networks before applying the ARACNE filtering algorithm^33^ we observed a similar trend for both enrichment and binary classification than with the filtered datasets, albeit with lower odds ratio and F1 scores (**Supplementary Figure 4**). Lastly, using O4 interactions to predict BA.1 epistatic interactions we measured the highest F1 score for the December 2012 dataset (0.35 [0.02-0.67]), again consistent with the rise of this variant at the same time (**Supplementary Figure 5**). This validation using experimentally determined epistatic interactions suggests that efficient computation of MI values could be used to implement a near real-time surveillance system for epistatic interactions.

In order to put this idea of a near real-time surveillance system for epistatic interactions to the test, we focused on determining how early known epistatic interactions would be highlighted by the MI-based method. An ideal interaction pair is between positions 498 and 501 in the Spike protein, which has been shown experimentally to increase the Spike’s affinity to the ACE2 receptor both compared to the wild type and the single mutants alone, and thus the virus’ infectivity^20,34^. While mutations at position 501 were already observed in the Alpha variant, the double mutation was not observed before the Omicron variant appeared around November 2021 (Figure 4A). Consistent with this observation, we first predicted an O3 interaction between these two positions in the November 2021 dataset, which contained only 6 viral sequences out of 2,500 that were annotated as lineage 21K (Omicron BA.1, Figure 4B and D). In the following month the frequency of Omicron viral sequences already increased to 56.4% and as expected the interaction strength between the two positions increased to reach the O4 level. We observed that this interaction faded with time and eventually disappeared in February 2023, consistent with mutations having reached fixation in the population and thus bearing no further mutual information between them. While the time point datasets are cumulative and thus contain all sequences up until the focal time point, our sequence age based correction effectively eliminates the contribution of sequences older than a year in the calculation of the MI values (**Supplementary Figure 2**). This seems a desirable property for a real-time surveillance system that is focused on highlighting new epistatic interactions as they appear, even when the number of sequences bearing a double mutation is very low (*e.g.* N=6).

**Figure 4.**
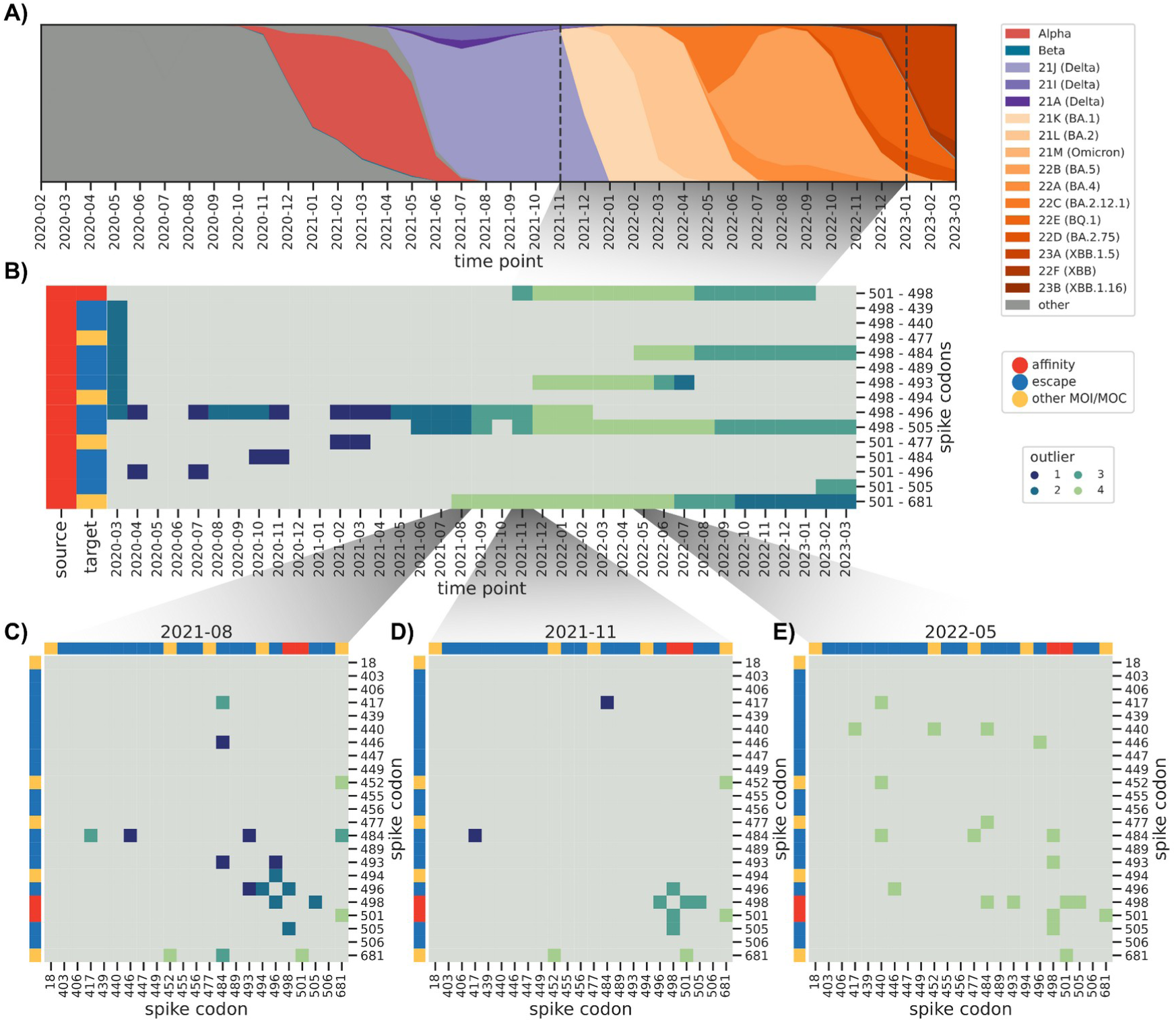
Predicted epistatic interaction in the Spike gene as a function of time. A) Muller plot indicating the relative frequency of each lineage as a function of time. B) Presence/absence matrix of predicted interactions between Spike codons 498/501 and those labeled either as escape variants or other Mutation of Interest/Concern. Gray indicates no predicted interaction. C-E) Interaction heatmaps between selected Spike gene codons at particular time points.

We observed a similar pattern for the interaction between either affinity related mutation (Spike codon 498 or 501) and escape or other mutations of concern, such as 501/484 (appeared for the first time in the Beta B.1.135 variant) already emerged in October 2020. In many cases the interactions were first estimated when the Omicron variant first appeared (around November/December 2021), consistent with the studies that have characterized mutations at these positions to interact epistatically and to modulate infectivity and immune escape^20,21^.

Another particularly interesting interaction we singled out was between spike codons 501 and 681, the latter having been described as conferring enhanced resistance to innate immunity^35^. Position 681 in particular has accumulated different amino acid substitutions in different variants: P681H in lineage 20I (Alpha B.1.1.7) and all Omicron subvariants, and P681R in lineage 21A (Delta B.1.617.2). Both positions have mutations that were first observed together when the Alpha variant emerged (around November 2020) and we therefore expected to see a predicted interaction with a high MI value. We however first estimated the 501/681 interaction in August 2021 (Figure 4C), which corresponded with the virtual extinction of the Alpha variant. We then checked the raw MI interactions, which contain indirect ones (**Methods**), and as expected the 501/681 interaction was first observed in November 2020 with outlier level O3, and which we observed every month until the last time point (**Supplementary Figure 6**). We posit that this apparent failure of the method to single out a known combination of mutations enhancing viral fitness was due to the relatively low number of sequences and their diversity until Summer 2021, which we have shown results in a large number of estimated interactions, many of which are likely false positives (Figure 3D).

The SARS-CoV-2 pandemic accelerated the usual pace of infection biology research: deep-mutational scanning and large scale antibody escape assays have been developed and released at unprecedented speed^36,37^. These are however necessarily limited to a single region of the genome of the virus: specifically the RBD region of the Spike gene. This necessarily excludes longer range interactions; not only within the Spike gene, as the 501/681 interaction, but also between different genes. Given the difficulty in testing those interactions in a laboratory assay, we focused on those for which we had the highest confidence to provide the community with a list of potentially interesting interactions. Overall the number of predicted interactions between genes is much lower than those within each gene, with the lowest number being 102 interactions in May 2022, and similar to the median number for the last 12 time points (137, Figure 5A). To focus on the highest-confidence predictions, we selected the inter-gene interactions that had one position in the Spike gene and had a large MI value (outlier level O4) in at least 9 time points; this resulted in 7 interactions (Figure 5B). Interestingly we observed three intergene interactions involving the notable 681 Spike codon, which is known to increase viral fitness. Among them we found codon 203 in the Nucleocapsid gene (N)^38^, which is known to increase infectivity. The other two positions with interactions with the 681 Spike codon were codon 26 in the ORF3a gene and codon 82 in the M protease gene; both are not known to influence viral fitness, and indeed they both are defining mutations for the Delta variant, their frequency closely following that of this variant (Figure 5C). We therefore suspect that these two interactions might be false positives, thus suggesting that despite the population structure correction, the method is sometimes still susceptible to the impact of the strong clonal interference observed with this virus and almost complete lack of recombination. It is however also possible that these mutations do have a yet to be determined impact on viral fitness.

**Figure 5.**
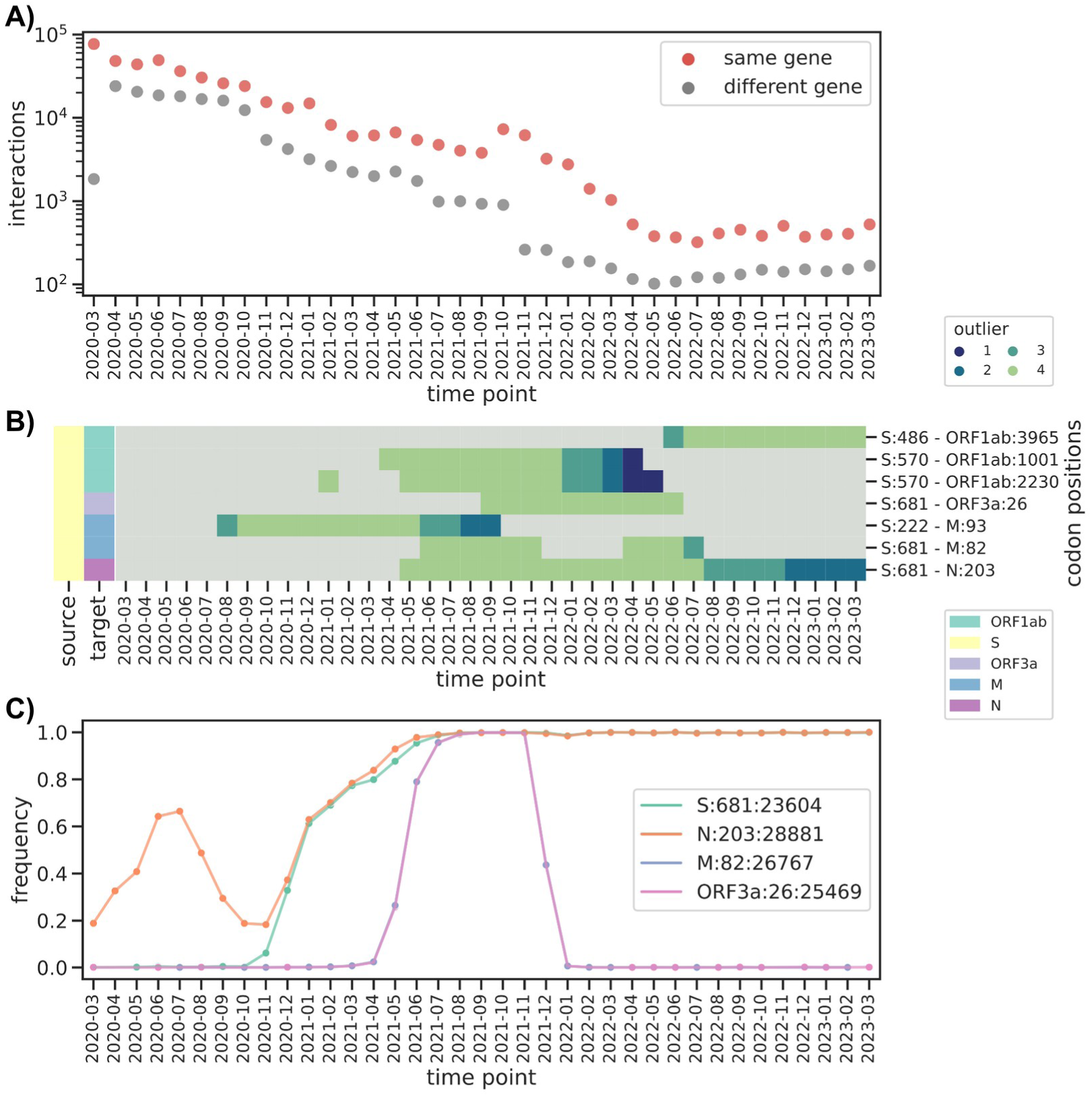
Between gene interactions. A) Change in within and between gene interactions across time points. B) Presence/absence matrix of predicted interactions between Spike codons and positions in other genes. Gray indicates no predicted interaction. C) Frequency of nucleotide substitutions at specific positions interacting with Spike codon 681. Labels are encoded with format gene:codon:nucleotide position. Purple and Pink lines overlap perfectly.

## Discussion

The tragic public health toll of the COVID-19 pandemic has been balanced by the unprecedented pace of scientific discovery and application in developing diagnostics and treatment solutions. Large scale sequencing of viral samples has played an important part in guiding public health interventions and more recently vaccine updates. The unprecedented scale of these genomic epidemiology efforts offer an ideal testbed for new computational approaches aimed at tackling future epidemics, for which genome sequencing will surely continue to play an important part^17^.

While multiple approaches to rapidly estimate the fitness effects of individual genetic variants have been proposed^13–15^, very few attempts have been made to estimate epistatic interactions between pairs of mutations. The interest in predicting them is not purely academic, as best exemplified by the appearance of the Omicron subvariants, each characterized by higher infectivity and immune evasion thanks to epistatic interactions, many of which have been experimentally verified^20,21^. Of note is the combination of amino acid changes at Spike codons 498 and 501, each of which alone results in a modest increase of the Spike protein for the ACE2 receptor, while together they have been shown to result in a >350 fold increase in ACE2 affinity, a clear example of positive epistasis. This large increase in fitness in turn likely allowed for the accumulation of slightly deleterious immune escape variants. Being able to quickly discover these interactions from the large number of viral sequences being routinely generated could complement predictors for single variants to form a reliable early warning system.

The few approaches that have so far been described in the literature make them relatively unsuitable for a near real-time system for predicting epistatic interactions. Neverov and colleagues recently proposed a method that relies on ancestral sequence reconstruction over a time-calibrated phylogeny to infer epistatic interactions based on mutations appearing one after the other over a relatively short time span^27^. This method is interesting in its similarity to what we presented here because it also explicitly includes a time component to aid prediction; it is however most certainly not able to scale to a large number of sequences and be efficient, as it relies on constructing a timed phylogeny and reconstruct the ancestral states for each branch of the tree. In this study we used a pseudolikelihood method, also termed direct coupling analysis (DCA) to validate the approach based on mutual information. Multiple studies have applied DCA to predict epistatic interactions in SARS-CoV-2, although at different times in the pandemic and with differences in the input alignments. Zeng and colleagues used DCA in the early stage of the pandemic (summer 2020) and reported a small number of predicted interactions, which is expected given the low diversity of the virus at that the time^39^. While DCA can be considered a more accurate model to predict epistasis, available implementations of the model tend to require more computational resources and more time to complete (∼4.5 hours on 100k Spike sequences using 16 cores), making them less suitable for a real-time system. One interesting exception is the study by Rodriguez-Rivas and colleagues, in which they used DCA over multiple sequence alignments spanning a longer evolutionary timescale^40^, which could be run as soon as the first SARS-CoV-2 genome sequence was available and required no subsequent update. Lastly, protein structure modeling has been used as a more direct way to measure the impact of combinations of mutations on the binding of the Spike protein to the ACE2 receptor^40^, a method that is likely unable to scale to thousands of sequences.

We chose to use a method based on computing mutual information between pairs of positions for its balance between speed and precision/sensitivity. We have in fact shown how reasonably accurate reconstruction of known interactions can be obtained with as little as 10,000 sequences with a short computation time (∼2 hours to process 10,000 sequences). These characteristics make it an attractive approach to be integrated into automated systems that feed on central sequence repositories. We introduced a sequence weighting strategy based on the age of each sequence so that a higher score would be given to emerging interactions and old ones be gradually removed. We showed how we were able to predict the known Spike 498/501 interaction as early as 6 sequences encoding the double mutations were present in the dataset, thus demonstrating its potential use as an early warning system.

One challenge of this genome-wide method is the interpretation of interactions predicted between genes^18^. The mechanism behind interactions within a single gene is intuitive; in the case of the Spike protein many immune escape mutations destabilize the protein structure while providing a fitness advantage. In more general terms, changes in amino acid sequences might result in protein structure changes that might need to be compensated by changes in residues that are close to each other in the protein structure. In case of an interaction between protein coding genes, unless these proteins share an interaction interface, two possible explanations are left to explain the prediction of an interaction: a functional interaction or a false positive. We found compelling that 3 between gene interactions out of 7 that we flagged as high quality involved the known 681 Spike codon, one of which with the known 203 codon in the Nucleocapsid gene, whose mutations result in higher infectivity^38^. Given the recent report on the importance of intersegment (i.e. between genetic backgrounds) epistatic interactions in modulating the evolution of the hemagglutinin gene in the flu virus^41^, we believe that these longer range interactions are worth reporting and explored in further detail.

With this study we have tested the applicability of a relatively simple computational method to estimate epistatic interactions from a large collection of viral sequences. By leveraging the metadata associated with each sample, notably the collection date, we could track the dynamics of these interactions, which can be used to filter out spurious predictions and prioritize other ones that appear over multiple subsequent time points. This approach is however heavily reliant on the quality of the metadata; as we have shown we could identify the interaction between the 498/501 spike codons with as little as 6 samples bearing the double mutation. During our earlier analysis we noticed how we could predict this interaction even earlier (December 2020, **Supplementary Figure 7**), which puzzled us because at the time the Alpha variant, which only had mutations at the 501 codon was increasing in frequency. Upon careful inspection we discovered that 2 Omicron sequences out of the 2,500 used in the December 2020 timepoint were erroneously dated, thus generating a modest signal. The analysis presented in the main text has been performed after a further round of filtering out sequences that are known to have incorrect metadata. This accident speaks in favor of the high sensitivity of the method, but also for the need of a well-curated sequence repository, which requires proper funding, infrastructure and a community engaged in curating it^17^. The potential of such repositories in enabling real time surveillance and interventions in the face of rapidly unfolding epidemics are well worth the effort.

## Materials and Methods

### Dataset and estimation of epistatic interactions

We used a mutual information (MI) based method to estimate epistasis across multiple sequence alignments (MSAs) of SARS-CoV-2 sequences. This method is based on a previously developed software (spydrpick^31^). Our implementation, derived from that provided in the panaroo software^42^, was optimized to process a very high number of sequences in a short amount of time.

We selected the whole pan-genome alignment available on the GISAID platform, counting 15,372,583 sequences by mid April 2023. We only extracted the high quality sequences according to a curated GISAID tree^43^ and which were also available in the public NCBI database (N=5,447,247). Also, since the non-coding 5’ and 3’ ends of the SARS-CoV-2 genomes are generally of lower quality, we resized them, considering only the positions from 266 to 29,768 (98.7% of the alignment length). Sequences were then deduplicated (N=4,093,019) and stored in a matrix file. Before computing the mutual information between positions, sequences were weighted according to their distance to the root of the phylogenetic tree, normalized by the number of leaves at each internal node. This in turn ensures that the contribution of a large number of very similar sequences to the final prediction is reduced.

The actual MI value was computed in the same way as the original implementation^31^. Shortly, mutual information is computed between each pair of residues in a multiple sequence alignment, which can take one of 5 values for each sample, the four nucleotides plus a character for gaps or undetermined bases. The main equation is as in the original implementation, and as follows:

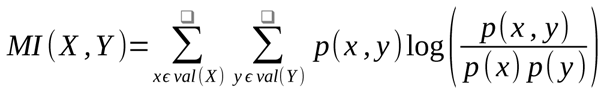

in which *X* and *Y* indicate the two positions in the multiple sequence alignment, *val* (*X*) and *val* (*Y*) the five possible discrete states at both positions, *p*(*x, y*) the joint probability of *X* =*x* and *Y* = *y*, and *p*(*x*) and *p*(*y*) the marginal probabilities. As described in the original publication, the joint probability *ṗ*(*x, y*) has to be estimated from the data itself, with the following equation:

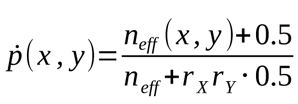

in which *n_eff_* (*x, y*) indicates the effective count of occurrences of the *x* and *y* pair (see below), *n* the number of samples, and *r _X_*=*val* (*X*), *r_Y_* =*val* (*Y*). Marginal probabilities are computed with equations:

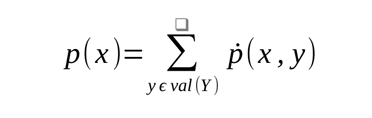

and

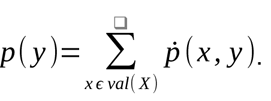

As indicated above, we introduced a sample weighting strategy similar to that used in the original implementation, and based on the phylogenetic tree of all samples. For each sample *i*, we computed the weight *w_i_* as the distance from the root, with a normalization step such that the length of each branch is normalized by the number of leaves downstream of each branch:

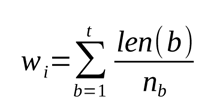

in which *b* is the internal branch over the total number *t*, *len* (*b*) the length of the branch, and *n_b_* the number of downstream leaves. The weights are used to derive *n_eff_* (*x, y*), which is the count of of occurrences of the *x* and *y* pair multiplied by the weights.

We ran our implementation of the spydrpick algorithm on a random sample of 100 positions in the MSA to extract a MI threshold based on the 90th percentile of the computed MI values. The resulting interactions were then filtered by an algorithm implemented for inferring gene expression networks (ARACNE^33^), and used to retain direct interactions, discarding indirect ones with a lower MI value. The output is generated as a tab-separated values (tsv) file. We defined 4 thresholds to indicate our confidence in the computed MI values based on the Tukey method as follows:

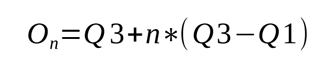

Q3 and Q1 respectively indicate the upper and lower quartiles of the MI values, and *n* are four different coefficients (*n*={1.5, 3,6, 12}) to be multiplied to the interquartile range, one for each outlier level. The resulting output with MI values was then annotated using the SARS-CoV2 GFF file (RefSeq NC_045512.2) and each pair of positions were associated to the respective gene, codon number, and gene relative codon number. We excluded interactions within the same codon and between adjacent codons, as well as interactions between different genes whose nucleotide distance was < 2. For the overall analysis (Figure 1, **Supplementary Table 1**) we ran the pipeline on the whole dataset of public sequences up to March 2023. A total number of 4,093,019 deduplicated and weighted sequences were processed. Smaller subsets of 1,000, 10,000, 100,000, and 1,000,000 sequences were generated by drawing random sequences from the complete dataset.

### Validation of predictions

We validated the interactions using a set of experimentally validated epistatic interactions and notable positions in the RBD of the Spike protein. We divided them into three sets, based on their known impact on viral fitness; we termed codons 498 and 501 as “Affinity” mutations for their positive epistatic interaction resulting in increased affinity to the ACE2 receptor^21^. We termed codons 406, 417, 446, 447, 449, 484, 493, 496, 505, and 506 as “Escape” mutations for their contribution to immune escape, especially in the 498/501 genetic background^20,21^. We added to this category those Spike codons with a relatively high escape score (> 0.1) as computed by the Escape Calculator^44^. Lastly, we added other notable Spike codons based on their designation as Mutation Of Interest or Mutations Of Concern if they were not already included in the other two sets: 18, 439, 452, 477, 494, and 681^32^.

We used interactions between the codons in all three sets if they were in the RBD region (318 < codon < 541) and three different methods to validate our predicted interactions. The first method is based on an hypergeometric test (Fisher’s test) for the enrichment of observed known interactions over all possible RBD interactions. We defined the universe of all possible interactions as those between residues mutated in at least one sequence of those used to build the predictions. We then compared the odds-ratio thus computed against that from 1,000 permutations of the actual RBD network, avoiding self links (a position interacting with itself) and those between adjacent codons. For the second validation method we built a binary classifier to indicate whether the interactions passing the first MI value threshold (90th percentile of MI values computed over 100 random positions) were the known ones or not. For each outlier level (O1, O2, O3, O4) we used its threshold value to classify the interactions and computed the F1 score using the known interactions as a truth set. For the third validation we used the inferred pairwise epistatic coefficients between 15 BA.1 mutations and tested the ability of our predicted interactions to predict interactions with a coefficient > 0.4, using the outlier levels as thresholds. For all three validation methods we measured the 95% confidence interval of each indicator (i.e. odds-ratio, specificity, sensitivity, F1 score) through bootstrapping with N=1,000.

### Time-resolved analysis

We also tested the accuracy of our method on different time-points along a roughly three year period (December 2019 to March 2023). To this purpose, we divided our dataset in different subsets of sequences for every month as follows. Firstly, we binned the sequences for each month, collapsing the period between December 2019 and February 2020, since the number of sequences collected in the databases was very low in the very beginning of the pandemic (N=592), obtaining a total of 38 bins. Furthermore, we randomly selected 2,500 sequences for each month filtering out those known to have an erroneous sample collection date^45^. For each subset we then created a MSA with the selected sequences at that specific time point plus all the previously selected sequences. Moreover, beyond the phylogenetic weights, a second time-based weighting system was applied to the sequences in order to maximize the importance of emerging interactions. This has been made to reduce the MI values of less recent interactions. For this purpose we developed a function of exponential decay that follows a Hill curve function for the weighting of each sequence. We used a Hill coefficient of 3 and a weight of 0.5 at 120 days (**Supplementary Figure 2**) . Both phylogenetic and time-based weights were then combined via multiplication to generate a final weight *w_i_* for each sequence in each subset. We used Nextclade^46^ v2.14.0 to generate the list of amino acid substitutions in each sequence as well as their lineage, and used the pyfish package^47^ v1.0.3 to generate a Muller plot to represent the changes in lineage relative proportions across all subsets.

### Comparison with direct coupling analysis (DCA)

To further validate our data, we also used a pseudolikelihood method called DCA (Direct Coupling Analysis), and implemented in the plmc tool^24^ that calculates the covariation and coevolution of biological sequences by inferring undirected graphical models, and applies a pseudo-likelihood approximation (Potts model) to impute the interaction strength between all pairwise positions in a given sequence. Since this implementation scales poorly with large MSA, both in terms of length and depth (*i.e.* number of sequences), we used the spike portion of the MSA in the last time-resolved dataset (i.e. all selected sequences until March 2023) to compare values computed using plmc against MI values. We used the following command line arguments when running plmc (commit 18c9e55): “--fast -m 20 -le 20.0 -lh 0.01 -a -AGCT”. To have a more direct comparison we did not apply the time-based weighting for MI values but only that based on phylogenetic distances.

### Code and data availability

All code used to compute MI values and run all the other analysis is available in the following code repository released under a permissive open-source license (MIT): https://github.com/microbial-pangenomes-lab/2022_sarscov2_epistasis. The code is based on bash scripts running a series of python scripts, using the following libraries: numpy^48^ v1.24.4, scipy^49^ v1.11.1, pandas^50^ v2.0.3, numba^51^ v0.57.1, treeswift^52^ v1.11.37, dendropy^53^ v4.6.1, networkx^54^ v3.1, scikit-learn^55^ v1.3.0, matplotlib^56^ v3.7.2, seaborn^57^ v0.12.2, and jupyter-lab^58^ v4.0.4.

We downloaded the input MSA and relative metadata and phylogenetic tree from the GISAID database^43^ on April 14th. We have restricted our analysis strictly to those sequences that were also deposited in public databases, as indicated at the following URL: https://hgwdev.gi.ucsc.edu/~angie/epiToPublicAndDate.latest. As such we believe we have respected the stringent end user agreement imposed by GISAID.

## Author contributions

MG conceived the study, GI and MG wrote the analysis code, performed the analysis, assembled figures and wrote the manuscript. Both authors have no conflict of interest to disclose.

## Acknowledgements

We gratefully acknowledge all data contributors, i.e., the Authors and their Originating laboratories responsible for obtaining the specimens, and their Submitting laboratories for generating the genetic sequence and metadata and sharing via the GISAID Initiative, on which this research is based. We thank Theo Sanderson and Chris Lauber for their feedback on the manuscript draft. MG and GI were funded by the Deutsche Forschungsgemeinschaft (DFG, German Research Foundation) under Germany’s Excellence Strategy - EXC 2155 - project number 390874280. GI was further supported by a FEMS Research and Training Grant (FEMS-GO-2020-258). Funders had no role in study design and in presentation of the results.

## Supplementary Material

**Supplementary Figure 1.**
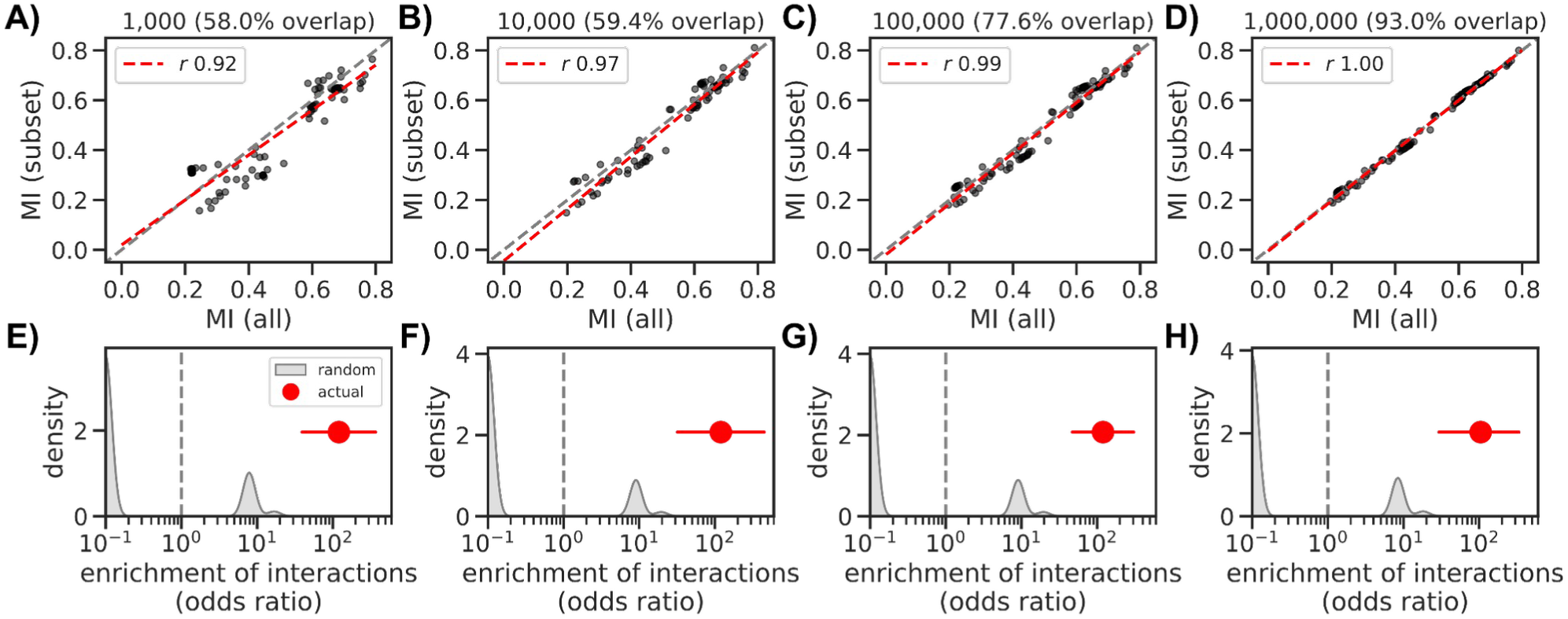
Influence of dataset size on mutual information estimation, using only O3 and O4 interactions. (A-D) Linear regression analysis of the intersection of mutual information values between interactions belonging to Outlier level 3 and 4 in the whole dataset versus the same Outliers in reduced datasets of (A) 1000, (B) 10,000, (C) 100,000 and (D) 1,000,000 sequence subsets. (E-H) Enrichment of interactions between Outliers 3 and 4 positions known to epistatically interact (vertical red line) versus a series (N=1,000) of random RBD networks with the same number of interactions as the real one (gray distribution). Subsets are the same as the panel directly above: 1,000 (F), 10,000 (G), 100,000 (H), and 1,000,000 (I).

**Supplementary Figure 2:**
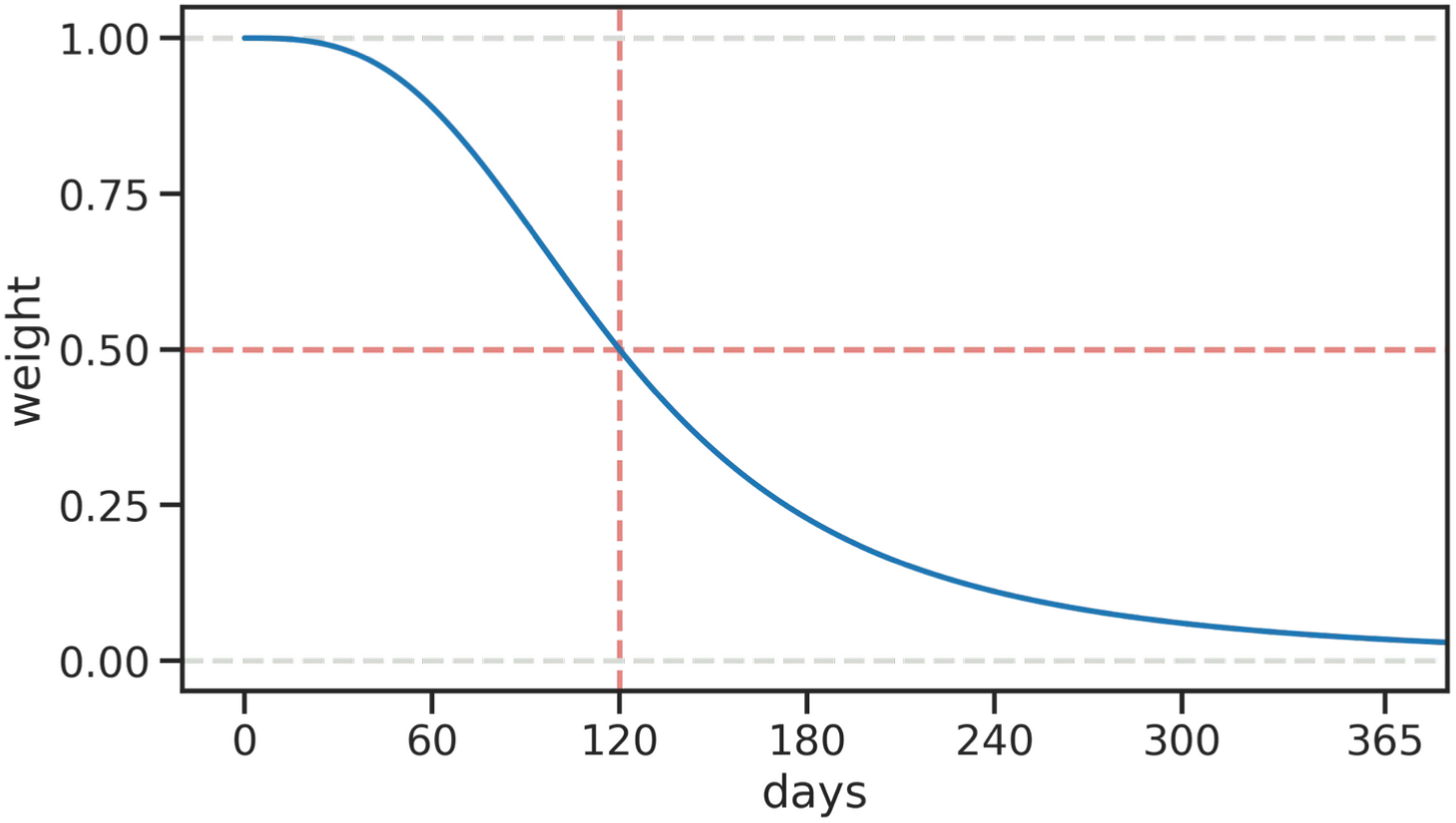
Representation of the Hill curve function that was used for the time weighting. The Hill coefficient was set to 3, and the function allowed a reduction of 0.5 of the initial sequence weight (weight=1 at the focal time point) after 120 days (∼4 months).

**Supplementary Figure 3.**
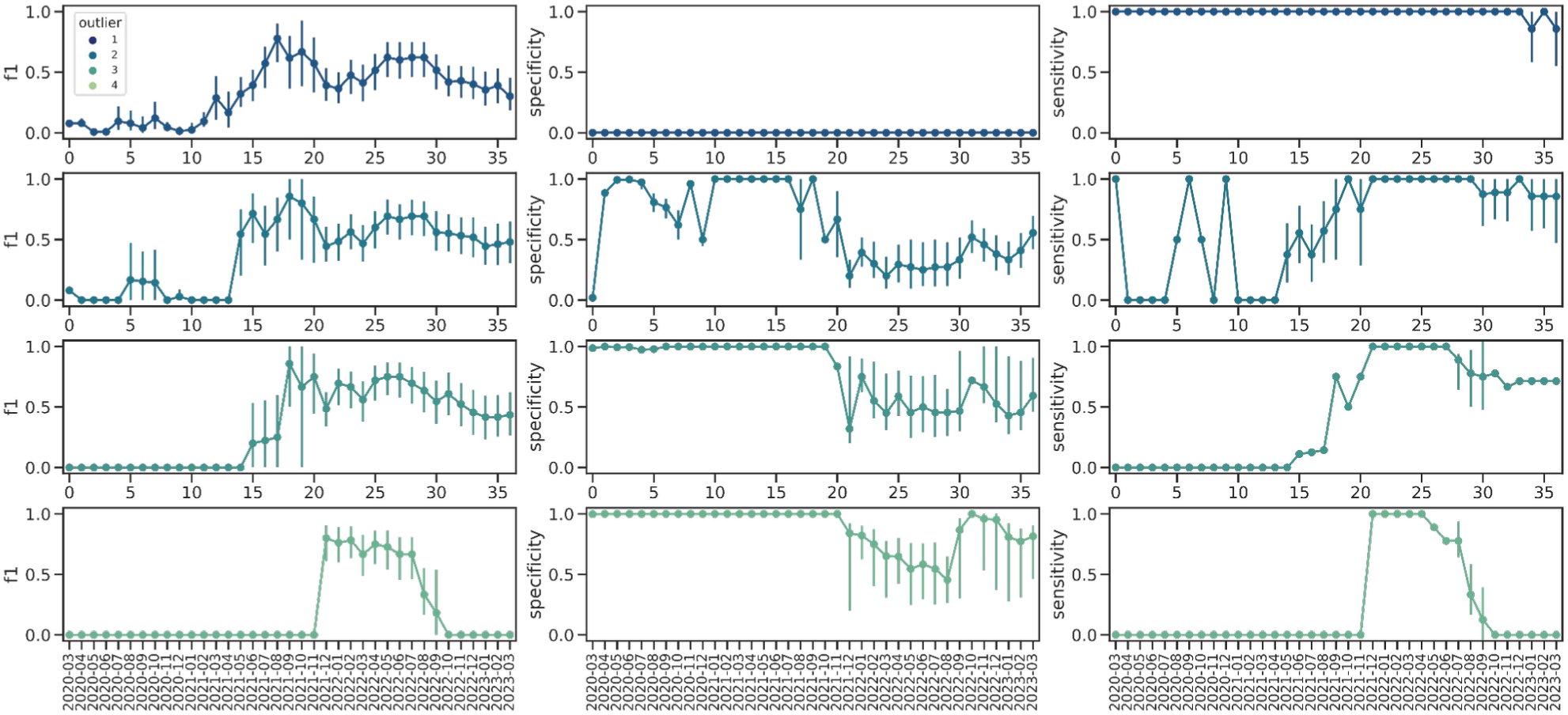
Alternative validation using known RBD epistatic interactions. Performance of a binary classifier for RBD interactions using the four outlier thresholds (rows) and measuring three different indicators: F1 score (first column), specificity (second column), and sensitivity (third column). Vertical solid lines indicate the 95% confidence interval.

**Supplementary Figure 4.**
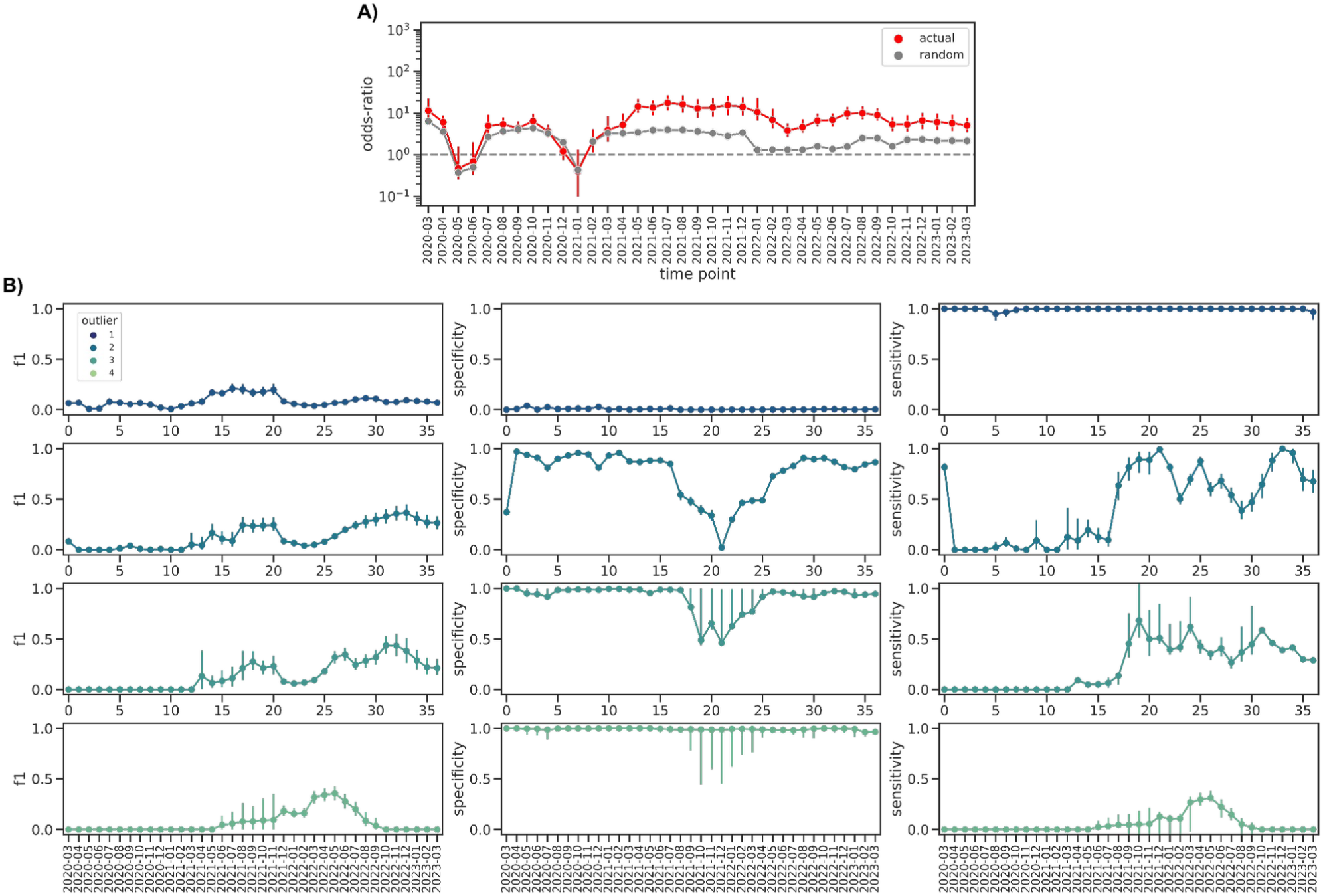
Validation using predicted interactions before filtering through the ARACNE algorithm. A) Enrichment of known interactions for the actual predictions (red) and for 1,000 random interaction networks (grey). Vertical red lines indicate the 95% confidence interval, grey shaded area the standard deviation. B) Performance of a binary classifier for RBD interactions using the four outlier thresholds (rows) and measuring three different indicators: F1 score (first column), specificity (second column), and sensitivity (third column). Vertical solid lines indicate the 95% confidence interval.

**Supplementary Figure 5.**
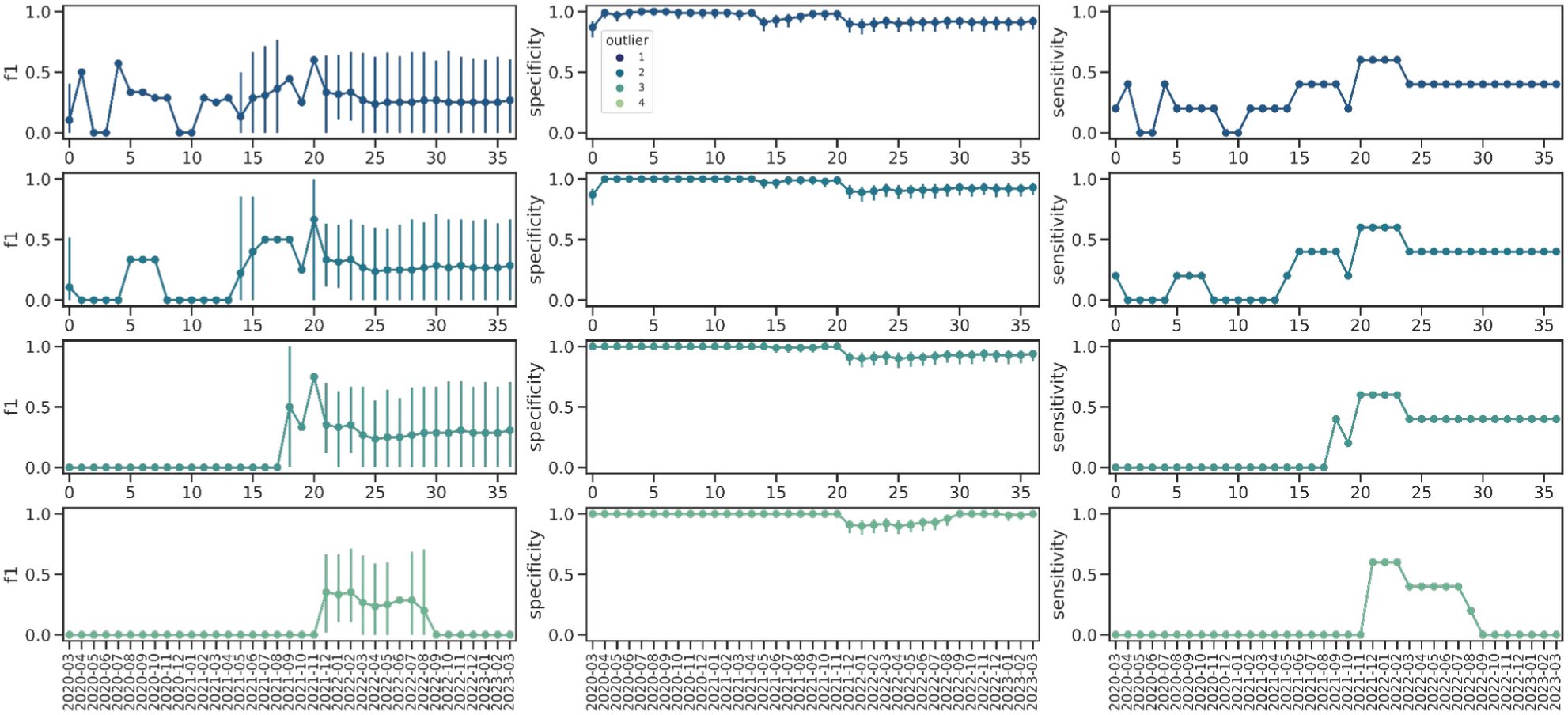
Alternative validation using pairwise coefficients for 15 BA.1 mutations. Performance of a binary classifier for RBD interactions using the four outlier thresholds (rows) and measuring three different indicators: F1 score (first column), specificity (second column), and sensitivity (third column). Vertical solid lines indicate the 95% confidence interval.

**Supplementary Figure 6:**
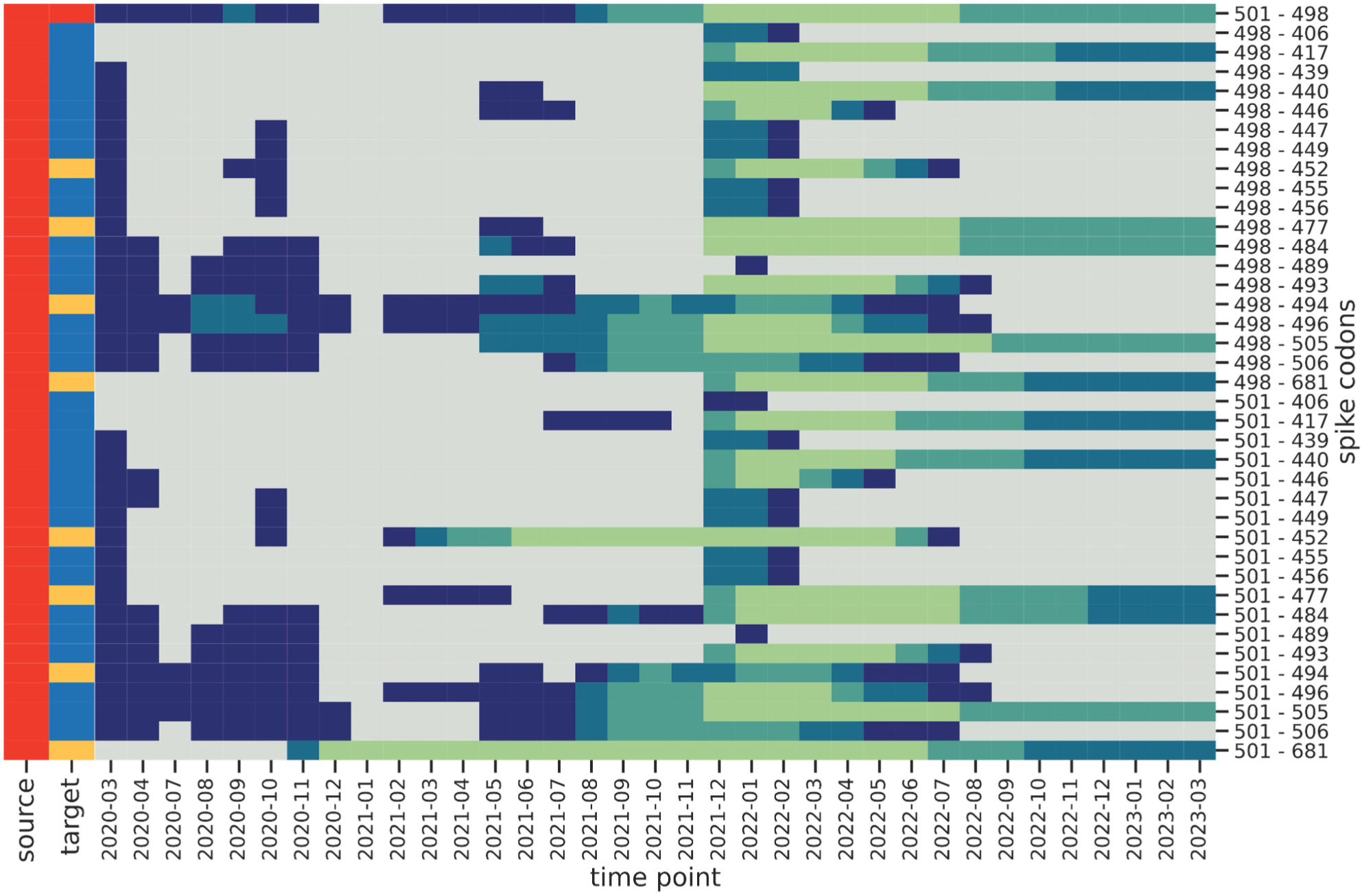
Presence/Absence matrix of notable spike positions using the raw MI interaction data before the filtering with the ARACNE algorithm. Gray indicates absence of predicted interactions.

**Supplementary Figure 7.**
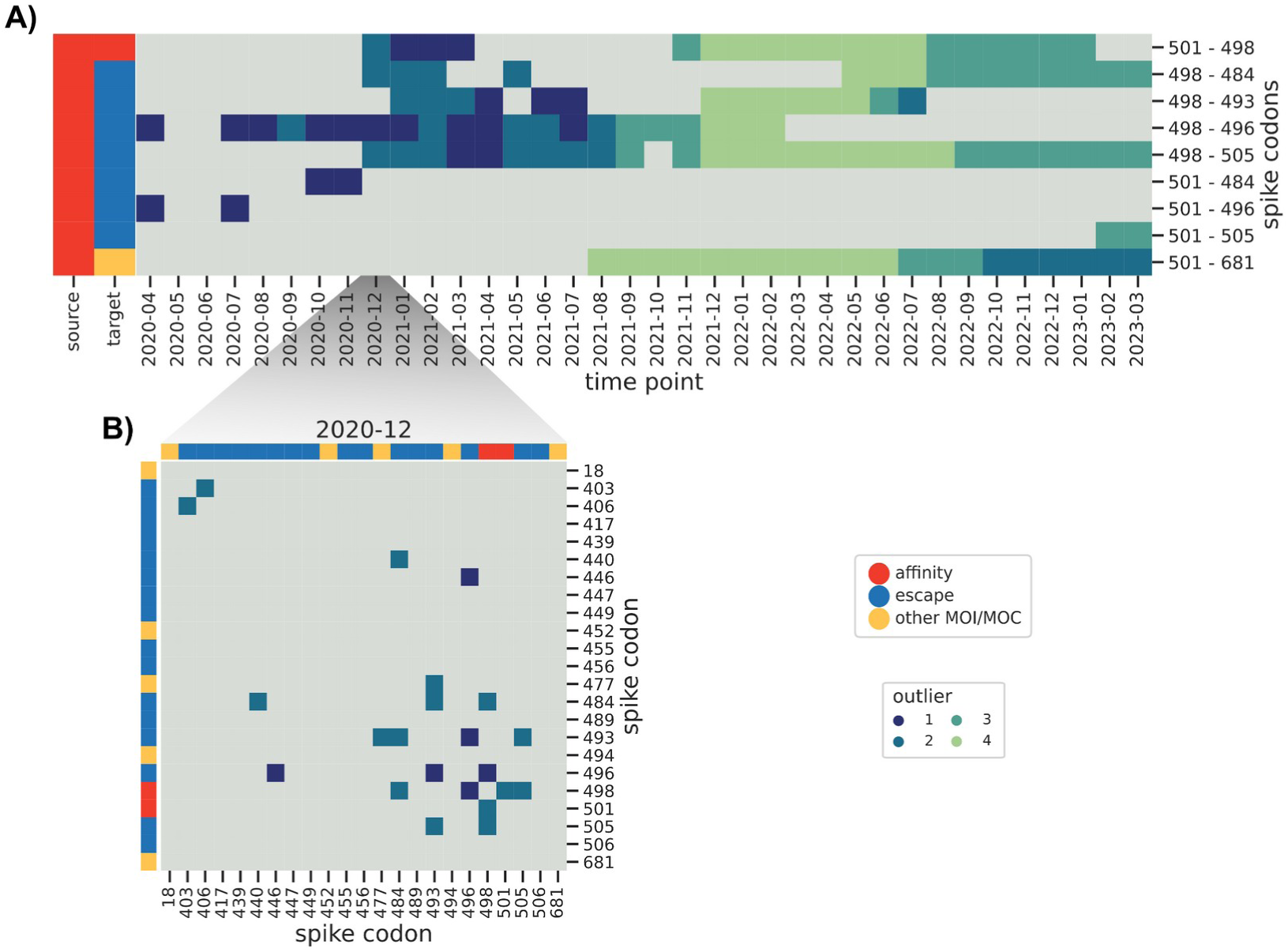
Predicted epistatic interaction in the Spike gene as a function of time, before the removal of sequences with incorrect dating. A) Presence/absence matrix of predicted interactions between Spike codons 498/501 and those labeled either as escape variants or other Mutation of Interest/Concern. Gray indicates no predicted interaction. B) Interaction heatmaps between selected Spike gene codons for December 2020.

**Supplementary Table 1:** predicted interactions from the complete dataset.

**Supplementary Table 2:** predicted interactions from the March 2023 dataset, without the time weighting correction.

**Supplementary Table 3:** enrichment of known interactions in each time subset. For each month between March 2020 and March 2023 the enrichment value for the actual dataset is reported, as well as the average values for 1,000 randomized RBD interaction networks.

**Supplementary Material 1:** predicted interactions for the subsets with sizes 1,000, 10,000, 100,000, and 1,000,000; columns are the same as those in **Supplementary Table 1**.

**Supplementary Material 2:** predicted interactions for the spike gene using the plmc implementation of the pseudolikelihood method; positions are relative to the first base of the spike codon, and all pairwise interactions are scored.

**Supplementary Material 3:** predicted interactions for the time subsets, from March 2020 until March 2023; columns are the same as those in **Supplementary Table 2**.

